# Reconstructing the developmental history of the cortex from postmortem tissue

**DOI:** 10.64898/2026.07.21.739857

**Authors:** Tatsuya C. Murakami, Nathaniel Heintz

**Affiliations:** Laboratory of Molecular Biology, The Rockefeller University, 1230 York Avenue, New York, NY 10065, USA

## Abstract

Single-cell-resolution mapping of the human brain is central to understanding the link between cellular-level phenotypes and disease. Tissue clearing and mesoscale imaging have facilitated organ-wide quantification of cell populations and are expected to drive the identification of pathological deficits in neurological disorders. However, mesoscopic comparison of human brain tissue remains challenging because of large variability in cortical folding patterns and heterogeneous cell distribution. Here, we present a “retrodictive” analysis of the cortex that reconstructs the proliferation history of neural stem cells from a static snapshot of tissue, providing a developmentally grounded coordinate system for comparison across samples. The analysis showed that most of the heterogeneity in cell distribution can be explained by the physical displacement caused by folding, which allows folding-induced variation to be removed and additionally reveals a possible mechanism of cortical folding. Together with a Bayesian framework, we estimated, with quantified uncertainty, the proliferation rate of a representative stem cell that summarizes the population of each cortical region. This analysis enables researchers to distill inter-individual differences down to stem cell properties and their distributions on the protomap. We anticipate our workflow to be a foundational framework for 3D neuropathological analysis, providing a common coordinate for the cortex-wide analysis of cells across individuals in development, aging, and disease.

## INTRODUCTION

Across evolutionary history, humans developed a markedly expanded cerebral cortex characterized by extensive convolution. The cortex carries out higher-order information processing such as cognition and voluntary movement, and harbors a diversity of cell populations. Untangling this cellular complexity is a central endeavor in neuroscience, not least because disruptions to cortical cell populations underlie neurological disorders (Braak and Braak, 1991; Pressl et al., 2024). Since quantitatively characterizing disease-driven selective cell loss can directly inform the identification of therapeutic targets, the quantification of cell-type composition and spatial organization in the cortex has taken on growing importance. This trend has been further accelerated by the advent of spatial transcriptomics and whole-organ RNA histology (Kumar et al., 2021; Kanatani et al., 2024; Liu et al., 2024; Murakami et al., 2026), which enable the simultaneous profiling of gene expression and spatial organization at cellular resolution.

In pursuit of precise detection of disease impact in cortex, the methods of quantitative histology have evolved considerably. Conventional two-dimensional tissue sectioning, a technique that remains widespread today, is highly sensitive to the angle and position at which sections are cut. Such methodological variability compromises the precision of quantitative measurements, making it difficult to accurately capture the effects of disease and giving rise to conflicting reports in the literature. In response to these limitations, stereology was developed as a framework to achieve statistically principled quantification within the constraints of two-dimensional sections (West, 1999). Stereology employs systematic random sampling to derive unbiased estimates of three-dimensional (3D) quantities from two-dimensional sections. More recently, 3D imaging of human cortex enabled by tissue clearing has emerged as a powerful alternative, bypassing the statistical extrapolation inherent to stereology and enabling direct identification of cell positions within an intact spatial context (Hama et al., 2015; Murray et al., 2015; Lai et al., 2018; Morawski et al., 2018; Hildebrand et al., 2019; Inoue et al., 2019; Park et al., 2019; Ku et al., 2020; Zhao et al., 2020; Pesce et al., 2022; Schueth et al., 2023; Oakes-Klein et al., 2026). Despite the advent of 3D histology, a fundamental challenge persists in cortical cell quantification: cortical folding introduces substantial variability in local cell distribution within individual brains, making it difficult to isolate the inter-individual differences, which is attributable to disease. For example, cortical thickness varies by more than 20% depending on whether the region is located on a gyrus (ridge) or in a sulcus (groove) (Fischl and Dale, 2000). Because the choice of region of interest is often defined by the investigator without accounting for such folding-related variability, quantification of disease-associated differences is substantially influenced by how the region is delineated, compromising reproducibility. Independently of histology, the field of medical image analysis, led by magnetic resonance imaging, has developed methods that account for convoluted cortical geometry in automated quantification. Among the most widely used are cortical flattening and surface inflation (Essen and Drury, 1997; Fischl et al., 1999). These cortical unfolding techniques are highly effective for visualization but treat the cortex as a two-dimensional manifold, disregarding the dimension of cortical thickness. It is therefore not applicable to cellular-resolution 3D histology. Another common approach is image registration to a standardized coordinate space, in which inter-individual differences in brain size and cortical folding pattern are normalized. While such registration is also employed in 3D histology (Menegas et al., 2015; Renier et al., 2016; Tatsuki et al., 2016), it does not necessarily preserve cell-level spatial correspondence, especially when applied to the human cortex, where inter-individual variability in geometry is substantial. Although these medical image analysis methods are grounded in rigorous geometric mathematics, the spatial transformations they impose generally lack biological interpretability. Thus, a biologically grounded normalization framework is needed for 3D histology.

A key observation is that cortical folding is supposed to be a mechanical consequence of developmental growth (Richman et al., 1975; Van Essen, 1997; Tallinen et al., 2014, 2016), and that the laminar structure of the cortex is established along neural stem cells, namely radial glial (RG) cells (Rakic, 1972). Building on this, we propose a biologically grounded spatial transformation of the cerebral cortex as a 3D manifold. We extracted the developmental trajectories formed by RG cells and defined a coordinate transformation that virtually smooths cortical convolution. Using the transformed coordinate as the basis for analysis, we discovered principles governing laminar thickness and formulated a growth model of RG cells that explains these observations. The growth model is consistent with a previously reported theory of cortical folding and explains how canonical sulci consistently form at their characteristic locations. Using the fact that the cortical laminae are built in an inside-out manner, with tissue forming progressively from the basal to the outer surface (Angevine and Sidman, 1961; Rakic, 1972), we examined the retrodiction of RG cell proliferation rates from a postmortem snapshot of a cortical piece. Using these rates as representative features of the observed brain regions, the analysis enabled precise detection of population changes across cortical regions. Beyond regional comparison, the framework can also be used to trace developmental, pathological, and evolutionary processes, offering a tool for deeper understanding of the cerebral cortex, the defining feature of the human brain.

## RESULTS

### Radial glial cell-guided biologically interpretable coordinate transformation

The analysis workflow for 3D histology in smooth-surfaced (lissencephalic) brains has been established around image registration. After whole-brain image acquisition, the image is typically registered to a standard atlas to obtain cell numbers per functional brain area (Menegas et al., 2015; Renier et al., 2016; Tatsuki et al., 2016). Cell populations are aggregated within pre-annotated areas, followed by statistical analysis. This workflow rests on an implicit assumption: that image registration provides a correct one-to-one correspondence between two individuals, and that within-region variability is not large (**Figure 1A**). However, this strategy breaks down when applied to a folded (gyrencephalic) brain that lacks a stereotyped folding pattern. Due to heterogeneous folding anatomy, there is no clear correspondence of a given point between two individuals, and high folding-induced within-region variability further confounds quantification.

**Figure 1.**
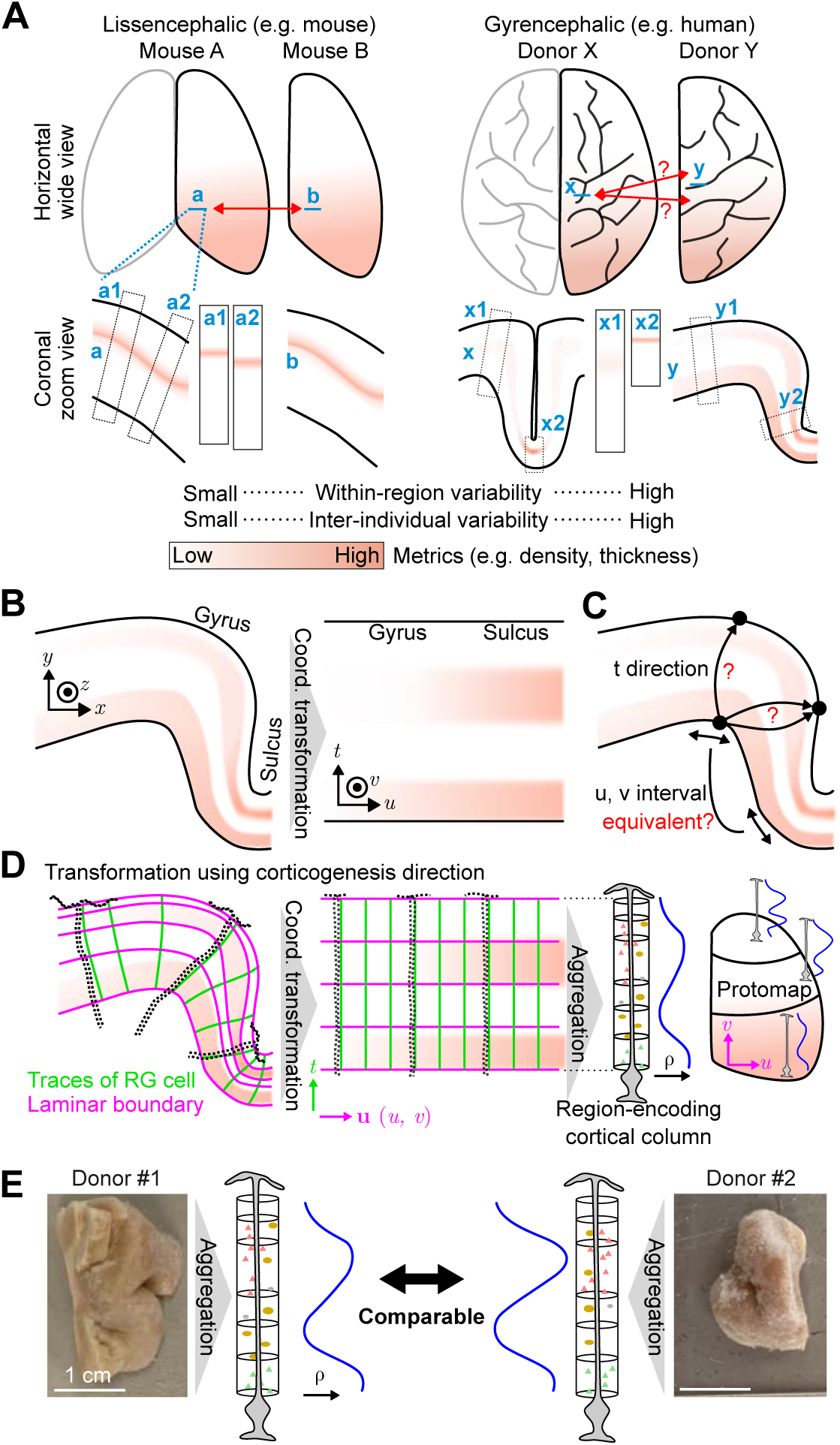
Challenges in the multiple-individual comparison of human cortex and proposed solution. (**A**) Illustration of the challenges in multiple-brain comparison in gyrencephalic brains that lack a stereotyped folding pattern. (Left) Comparison of lissencephalic brains is straightforward and often achieved by image registration. A position in one brain can be readily identified in the other. This also holds for brains with a stereotyped folding pattern. (Right) In highly convoluted gyrencephalic brains such as the human, no obvious corresponding point exists between two brains. In cell-scale analysis such as light-sheet imaging, this problem becomes non-negligible. The fact that cortical thickness varies with position along a sulcus further confounds cell number quantification. (**B**) Conceptual illustration of cortical coordinate transformation. (**C**) Challenge in coordinate transformation. The coordinate transformation cannot be uniquely determined from the surface positions of the cortex alone. (**D**) Illustration of the proposed coordinate transformation using the traces of radial glial (RG) cells and laminar positions. The transformed cortex can be readily reduced in dimensionality to a region-representing column. The idea is consistent with the protomap theory. *ρ* indicates the cell density profile of a ontogenetic column. (**E**) Illustration of multiple-individual comparison using the extracted region-representing columns.

We propose a framework that bypasses these limitations. Central to this framework is a biologically grounded coordinate transformation (**Figure 1B**). In the original image, coordinates are defined by three orthogonal Cartesian axes—*x*, *y*, and *z*—which are determined by the microscope system and carry no biological meaning. Our transformation redefines the coordinate system such that the same cortical layer lies on an identical tangential plane defined by two axes, *u* and *v*. This plane can be interpreted as a representation of the flattened cortical surface. The third axis, *t*, is defined to represent laminar position; for example, a given position within layer II is expressed by a specific value of *t* regardless of its *u*, *v* position. The transformation is diffeomorphic: a smooth, one-to-one, and reversible deformation that preserves topology. The challenge is that such a transformation cannot be uniquely determined from geometry alone. Given a position on the white matter surface *Swm*, there is no explicitly defined corresponding point on the pial surface *Spia*. Likewise, there is no explicit mapping between a unit increment in *u* and a distance in the original *xyz* space (**Figure 1C**). Previous approaches to this problem have relied on purely mathematical assumptions, such as Laplacian harmonic fields, to define this transformation (Jones et al., 2000). While geometrically valid, these transformations are not grounded in the underlying biology, which limits the interpretability of the resulting quantification. In contrast, we use a biological structure, specifically RG cells, to make the transformation unique. RG cells are neural stem cells that extend radially stretched fibers from the subventricular zone to the pial surface; cell division and migration occur along these fibers during corticogenesis (Rakic, 1972; Noctor et al., 2001). Using the trajectories of RG cells, we define the direction of the *t* axis. Using the spacing between RG cells, we define the intervals along *u* and *v*. The resulting *u*, *v*, *t* coordinate system has high biological interpretability. Cells surrounding a given *u*, *v* position correspond to cells that proliferated or migrated along the RG cell located at that position. This collection of cells can be regarded as a representation of an ontogenetic column, also related to a cortical column—a functional unit of the cortex generated by one or a few RG cells (Rakic, 1988). Thus, the quantification of within-region variability can be reframed as quantification of variability among ontogenetic columns, which is readily computed in the transformed coordinate system. Furthermore, if we can regress out the impact of cortical folding, we can obtain an aggregated, region-encoding column, effectively achieving dimensionality reduction (**Figure 1D**). The spatial arrangement of these region-encoding columns across the cortex constitutes what we term the protomap, connecting to Rakic’s protomap hypothesis, in which the prospective organization of cortical areas is established early in development at the level of progenitor cells (Rakic, 1988). This representation makes the quantification of inter-individual variability much more tractable and interpretable. This strategy is applicable even when only a portion of the cortex is available, as is often the case for diseased brains due to limited tissue availability, facilitating comparative neuropathology (**Figure 1E**).

### RG fibers, radial neurites, and penetrating arteries share a common radial orientation

Following the completion of neurogenesis, RG cells lose their ventricular attachments and transform into astrocytes (Voigt, 1989). The radial fibers of RG cells are therefore absent in the mature brain, precluding direct use for defining the coordinate transformation in adult tissue. Meanwhile, the cerebral cortex retains several structures that maintain the radial orientation established during development. Penetrating arteries descend perpendicularly from the cortical surface through the gray matter into the white matter (Duvernoy et al., 1981), and some neurites traverse the cortex in a similar radial fashion (Cajal, 1911). We hypothesized that these structures share a common radial orientation with RG fibers and could serve as practical proxies for defining the developmental orientation in the cortex.

To test this hypothesis, we first used the mouse brain, where both fetal and adult tissue are readily accessible, enabling direct comparison of RG fibers with candidate proxy structures across developmental stages. During embryogenesis, cortical blood vessels undergo remodeling, a process by which the vascular network acquires its mature architecture. Penetrating arteries have been reported to remodel from pre-existing vessels so as to align along RG fibers between E14.5 and E17.5 (Ma et al., 2013). To test whether this structural alignment holds across the entire cortex, we performed 3D imaging of the E16.5 mouse embryonic cortex using tissue clearing, staining blood vessels with lectin and RG cells with anti-BLBP antibody (Feng et al., 1994). We extracted directional vectors for each structure and quantified their angular correspondence. We found no significant difference in orientation between penetrating arteries and RG fibers (**Figure 2A**). Next, we examined whether this relationship extends to the cerebellum, which, unlike the mouse cerebral cortex, is folded. This is informative because the folded cerebellar cortex more closely approximates the geometry of the gyrencephalic human cerebral cortex. In contrast to cerebral RG cells, cerebellar RG cells (known as Bergmann glia) persist into adulthood. Penetrating arteries are also present in the cerebellar cortex. Using the paraflocculus of the adult mouse cerebellum, we co-stained blood vessels with lectin and Bergmann glia with anti-GFAP antibody, extracted directional vectors, and confirmed no significant difference in orientation between cerebellar penetrating arteries and Bergmann glia (**Figure 2B**). We next examined the alignment between penetrating arteries and neurites. Among neurites, we focused on the major trunks of apical dendrites and axons that run radially through the cortex, hereafter referred to collectively as radial neurites. We expect these radial neurites running along cortical columns. We used an adult Thy1-YFP-H mouse, in which YFP is strongly expressed in layer V pyramidal neurons, and co-stained the tissue for penetrating arteries with anti-αSMA antibody. We quantified their structural alignment and confirmed no significant difference in orientation (**Figure 2C**). Finally, we examined whether this alignment is conserved in the human cortex. We stained penetrating arteries with anti-αSMA antibody and myelinated fibers with anti-MBP antibody. Again, we confirmed no significant difference in orientation between these structures (**Figure 2D**). The alignment of penetrating arteries, radial neurites, and RG cells in the prenatal human brain has previously been suggested through immunostaining of tissue sections (Xu et al., 2014). Together with our findings, we conclude that the direction of either penetrating arteries or radial neurites can be used to reliably reconstruct the trajectory of RG fibers. Our data suggest that penetrating arteries and radial neurites may organize along RG fibers as a structural scaffold during development. This would be biologically parsimonious, as newly forming structures (penetrating arteries and radial neurites) are more likely to exploit an existing scaffold (RG fibers) than to establish an independent one.

**Figure 2.**
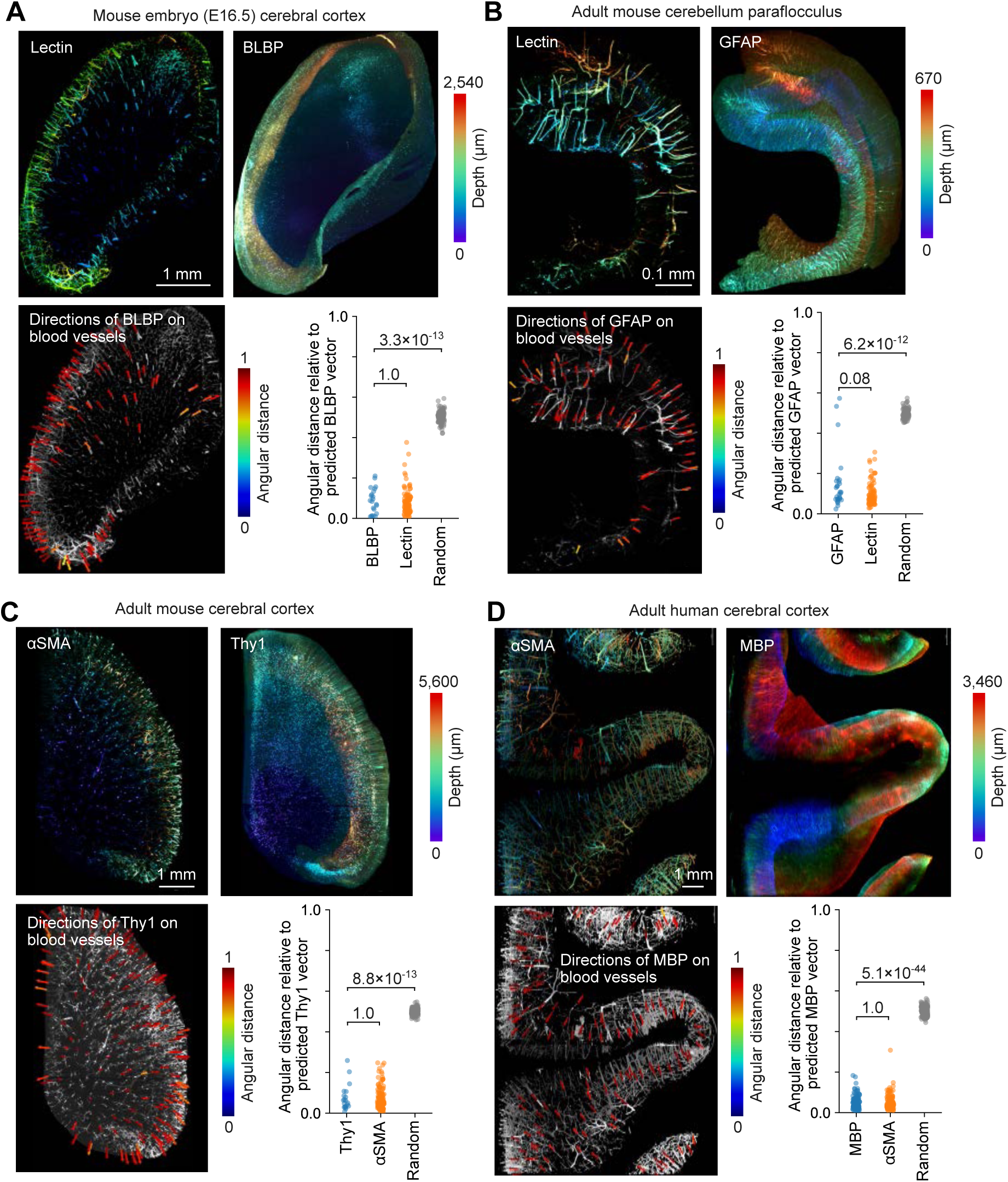
Co-aligned trajectory of RG fibers, penetrating arteries and layer-spanning neurites. (**A**) 3D rendering of an embryonic cerebral hemisphere showing lectin (top left, vasculature) and BLBP (top right, RG cells). Directions of RG cells overlaid on the lectin signal (bottom left) and quantification of the angular distance between these directional vectors (bottom right) are also shown. Random directional vectors were used as a control. (**B**) 3D rendering of a cerebellar paraflocculus showing lectin (top left, vasculature) and GFAP (top right, Bergmann glia). Directions of Bergmann glia overlaid on the lectin signal (bottom left) and quantification of the angular distance between these directional vectors (bottom right) are also shown. (**C**) 3D rendering of an adult cerebral hemisphere showing αSMA (top left, arteries) and Thy1 (top right, radial neurites). Directions of radial neurites overlaid on the αSMA signal (bottom left) and quantification of the angular distance between these directional vectors (bottom right) are also shown. (**D**) 3D rendering of an adult human cerebral cortex showing αSMA (top left, arteries) and MBP (top right, radial neurites). Directions of radial neurites overlaid on the αSMA signal (bottom left) and quantification of the angular distance between these directional vectors (bottom right) are also shown. The Mann–Whitney U test was used with Bonferroni correction for multiple comparisons.

### Morphogenic tracks as a representation of trajectory of RG fibers

Given that trajectories of penetrating arteries align with RG fibers, the directional course of cortical morphogenesis can be inferred from postmortem adult tissue, where RG cells are no longer present. Identifying the trajectories of these lost RG cells opens the possibility of inferring developmental processes, such as cell proliferation, directly from postmortem histology. We aimed to simulate these RG cell trajectories, which we term “morphogenic tracks”. If morphogenic tracks faithfully capture the developmental organization of the cortex, they should correspond to biological landmarks such as the boundaries between different functional regions of the cortex. We tested this expectation in a mouse brain.

We established an image analysis workflow for the reconstruction of morphogenic tracks from penetrating arteries. We construct a regression model that predicts the direction of morphogenic tracks as a continuous directional vector field **v**^ at any point in the cortex (**Figure 3A** and **Methods**). In addition to **v**^, reconstructing morphogenic tracks requires predefined starting points and a boundary at which the trajectory terminates. To this end, we placed seed points on the white matter surface and simulated their trajectories along the **v**^ until they reached the pial surface (**Figure 3B**). Using a mouse brain, we confirmed that the morphogenic tracks well reflected the trajectory of penetrating arteries (**Figure 3C**). Importantly, the morphogenic track was consistent with the functional boundary of brain areas obtained from Allen Brain Atlas (ABA) (**Figure 3D** and **Video S1**), indicating that the morphogenic tracks align with the functional parcellation of the cortex. Using a pig cingulate cortex, we also confirmed that our workflow can reconstruct the morphogenic tracks in a gyrencephalic brain (**Figure 3E** and **Video S2**).

**Figure 3.**
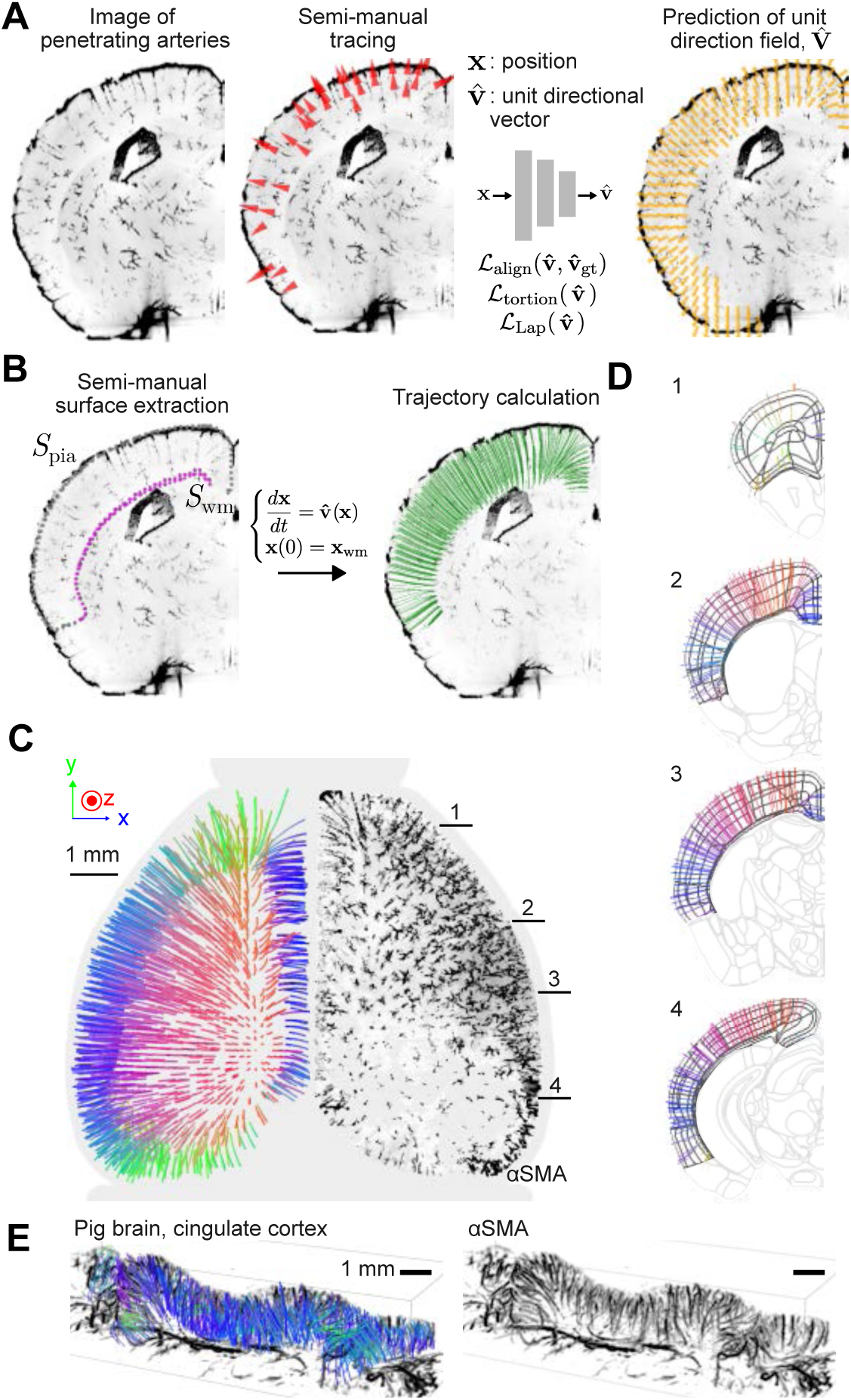
Morphogenic tracks. (**A**) Schematic illustration of the prediction of a unit-direction field representing the morphogenic flow from a 3D image of cerebral arteries (see also **Methods**). (**B**) Schematic illustration of the reconstruction of morphogenic tracks (see also **Methods**). (**C**) (Left) 3D rendering of morphogenic tracks in a mouse hemisphere. Color represents the directional components in *x*, *y*, and *z*. (Right) αSMA staining in the cortex. Only the staining within the cortical gray matter is shown. (**D**) Morphogenic tracks overlaid on the ABA functional boundaries. The coronal planes indicated in (**C**) are shown. (**E**) (Left) 3D rendering of morphogenic tracks in a pig cingulate cortex, overlaid with the αSMA signal. (Right) αSMA signal alone in the same region. The pial vasculature was removed for visibility.

### Morphogenic tracks as a basis for coordinate transformation

The vector field **v**^ is useful for simulating morphogenic tracks, but in its current form it is not directly suitable as a basis for coordinate transformation, as it retains only directional information. As discussed, an ideal coordinate transformation should be consistent with the process of corticogenesis and should assign the same *t* coordinate to a given cortical layer throughout the cortex (**Figure 1D**). This can be achieved by modifying the vector magnitude of **v**^ so that it reflects the stretch rate of RG fibers after cortical folding. The resulting modified vector field, termed **v**, then serves as the basis for the coordinate transformation along the radial direction *t* (**Figure 4A**). In the tangential direction, we expect that each point on the *uv* coordinate contains a uniform density of RG cells, or equivalently, of ontogenetic columns, making measurements at different *uv* positions directly comparable (**Figure 4B**). In other words, differences in measured values after coordinate transformation between two distinct locations can be attributed to genuine biological variation such as the proliferative capacity of RG cells. For clarity, we summarize the notation and meaning of our symbols in **Figure S1A**.

**Figure 4.**
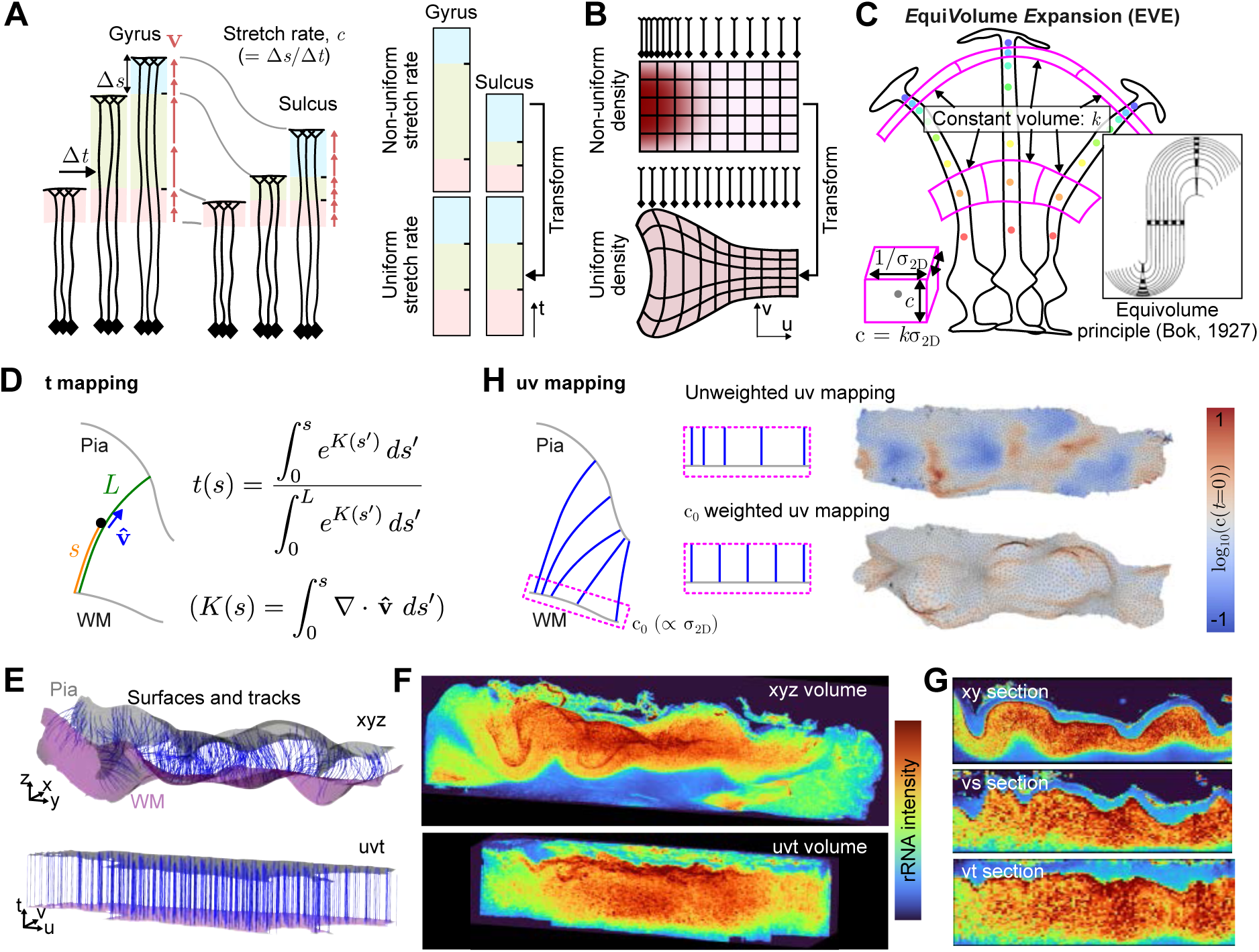
Transformation into morphogenic coordinate space. (**A**) Schematic illustration of the coordinate transformation in the radial direction according to **v**. The transformation is performed such that *Δs* represents the growth length during a time *Δt*. (**B**) Schematic illustration of the coordinate transformation in the tangential direction. The transformation is performed such that RG cells are distributed uniformly. (**C**) Illustration of the EVE growth model. Inset shows Bok’s original illustration. The magenta cube is a volume produced by a RG cell during a time interval *Δt*. The colors of the circles indicate *t*. (**D**) Implementation of the reparameterization of *s* to *t*. (**E**) *Swm*, *Spia*, and morphogenic tracks in *xyz* and *uvt* space. (**F**) Image of the rRNA-stained pig cingulate cortex before (top) and after (bottom) transformation into *uvt* space. (**G**) Cross sections of the original and transformed images. (**H**) Value-weighted *uv* mapping. *Swm*was mapped into *uv* space such that the predicted RG cell density is uniform. In **E**, **F**, and **G**, the scale of *t* was set to approximately match with that of *uv* for visualization purposes.

Although the realization of such an ideal coordinate transformation requires information about layer positions and the density of RG cells, these are not always available. We therefore explored whether an approximate coordinate transformation can be obtained using only geometrical information, *Swm*, *Spia*, and **v**^, treating the resulting morphogenic tracks as surrogates for RG fibers. To this end, we introduce a “*E*qui*V*olume *E*xpansion (EVE) growth model” that governs the rate of growth of morphogenic tracks, formulated as:

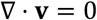

In the EVE growth model, all morphogenic tracks are assumed to be uniform, and each morphogenic track occupies a constant volume per unit *t* step (**Figure 4C**). The EVE growth model is grounded in the assumption that, within a given time span, each RG cell produces a similar total amount of cellular and extracellular material regardless of its position in the cortex, and that this volume is not distorted by cortical folding. The principal support for this assumption comes from Bok’s equivolume principle (Bok, 1929), a well-validated empirical rule which states that cortical layer segments preserve their volume across the folds of the cortex (Waehnert et al., 2014), whose biological basis has remained unknown. The EVE growth model inherently supports Bok’s equivolume principle and provides it with a neurodevelopmental basis.

We implemented the coordinate transformation according to the EVE growth model. Given that cortical formation proceeds in an inside-out manner from deep to superficial layers, we determined the direction of *t* such that at *t* = 0, the seed points of morphogenic tracks form the white matter surface, and at *t* = 1, the terminal points form the pial surface. Together with the EVE growth model, this boundary condition allows the *t* coordinate at a given position in the cortex to be determined analytically (**Figure 4D** and **Methods**). Next, we devised a *uv* coordinate mapping such that the density of morphogenic track is constant regardless of the position on *Swm*, while preserving the original surface shape as much as possible (**Methods**). As a demonstration, we imaged a cleared ribosomal RNA (rRNA)-stained cingulate cortex from a juvenile pig brain, approximately 35 mm × 10 mm × 10 mm in size, and used its gray matter for the coordinate transformation. The transformation was performed according to the morphogenic tracks (**Figure 4E**), as described above, and flattened the cortex as expected (**Figure 4F**). The laminae were more flattened than under the transformation without the EVE assumption, supporting our EVE assumption (**Figure 4G**). We confirmed that the density of the morphogenic tracks on the white matter surface were equalized after the transformation (**Figure 4H**).

### Precise laminar thickness prediction with morphogenic tracks with EVE growth model

Accurately predicting laminar position within the cortex has long been an important problem in medical imaging for precise cortical segmentation. We reasoned that quantifying cortical layer thickness, captured by the stretch rate *c* (= *∂ s*⁄*∂ t*, where *s* is radial arc length; see **Figure S1**), could reveal the physical principle governing cortical thickness. By flattening the folded cortex into a continuous space, our transformation lets us measure stretch rate of ontogenetic columns at every point, rather than reducing the analysis to a binary gyrus-versus-sulcus contrast. Because this representation is continuous, it permits analytical, calculus-based investigation of laminar formation and folding. We hypothesized that if cortical morphology is governed by simple physical laws as suggested by previous studies of cortical folding (Tallinen et al., 2014, 2016), then its internal structure, specifically laminar thickness, should exhibit relatively simple regularities.

If a perfectly flat cortex existed, the thickness of each cortical layer would be uniform, and the stretch rate *c* of ontogenetic column would be constant. In reality, the stretch rate varies with position. We examined how much the stretch rate deviates from the flat condition and whether this deviation exhibits any spatial pattern. Using laminar positions obtained from the rRNA signal as ground truth, we computationally flattened the pig cortex so that the same layer appears at the same *t* position (**Figure 5A** and **Methods**). We then obtained *Δc*, the difference between the actual stretch rate and the depth-invariant stretch rate that would occur in the flattened state. Analysis of *Δc* would provide a deeper understanding of how laminar thickness changes along tangential direction. To this end, we performed singular value decomposition (SVD) on *Δc* to extract dominant patterns explaining it. We found the rank-1 energy fraction to be 97.3%, indicating that laminar thickness is approximately determined by two separable factors: *α*(*u*, *v*), which depends only on the tangential direction, and *f*′(*t*), which depends only on the radial direction, with no interaction term depending on both. We confirmed that this rank-1 approximation reconstructs *Δc* (**Figure 5B**). The shape of *f*′(*t*) obtained through SVD motivated us to examine whether a simple linear function of *t* (t-linear function) is sufficient to achieve cortical flattening. Remarkably, a t-linear *f*′(*t*) was sufficient (**Figure 5C**). In summary, we found that the stretch rate can be formulated as:

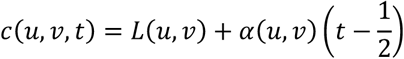

**Figure 5.**
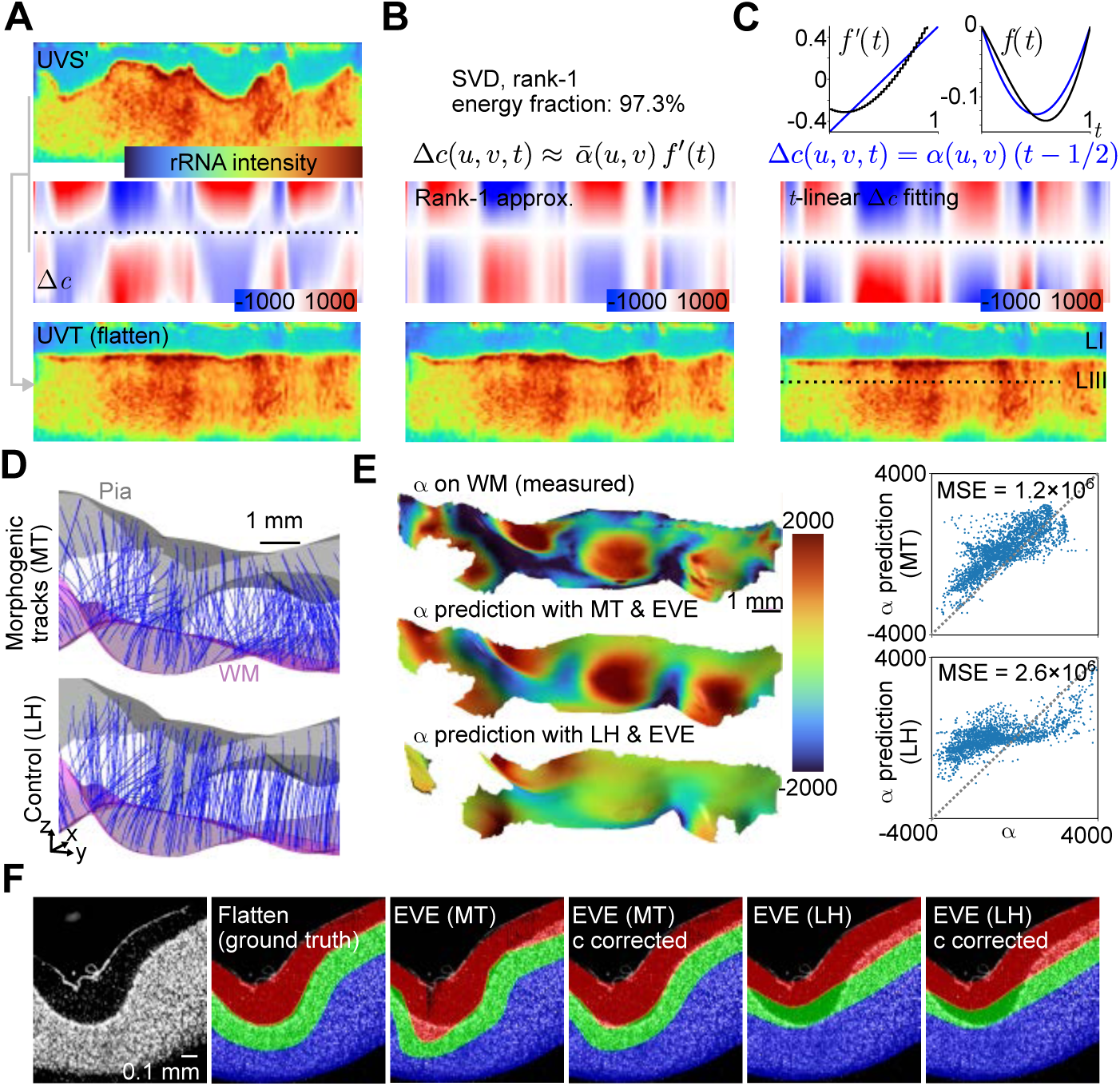
Precise laminar thickness prediction with morphogenic tracks with EVE. (**A**) (Top) rRNA intensity map in a representative section of the *uvs*′ coordinate. *s*′ is *s* linearly normalized by cortical thickness, such that *s*′ = 0 at *Swm* and *s*′ = 1 at *Spia*. (Middle) Difference in stretch rate, *Δc*, between the actual stretch rate and the stretch rate in *uvs*′. *Δc* encodes the displacement required to flatten the cortex. (Bottom) Representative section of the flattened cortex in the *uvt* coordinate after applying the displacement. (**B**) (Top) SVD rank-1 energy fraction of *Δc*. (Middle) Rank-1 approximation of *Δc*. (Bottom) Representative section of the flattened cortex after applying the displacement obtained from the rank-1 approximated *Δc*. (**C**) (Top left) *f*′(*t*) obtained from SVD. The line *t* − 1⁄2, used for the approximation of *f*′(*t*), is drawn for comparison in blue. (Top right) The integral of *f*′(*t*), which encodes *Δs*, and the integral of *t* − 1⁄2 in blue are shown. (Middle) *Δc* obtained under the assumption that *Δc* is linear with respect to t. (Bottom) Representative section of the flattened cortex after applying the displacement obtained from the t-linear *Δc*. (**D**) Morphogenic tracks obtained from the morphogenic vector field (top) and from a vector field generated by the Laplacian harmonic potential function (bottom). (**E**) Comparison of measured and predicted *α*, shown as heatmaps (left) and scatter plots (right). (**F**) Prediction of laminar position using *c* obtained from various prediction approaches.

Here, *α* is the stretch acceleration (*∂*^2^ *s*⁄*∂ t*^2^, the rate of change of stretch rate along *t*) and is constant with respect to *t* (t-constant). Because cortical thickness, *L*(*u*, *v*), is readily measured, this formula indicates that knowing the *α* alone is sufficient to determine laminar thickness (i.e. stretch rate *c*).

The next key question is whether *α* can be accurately predicted from cortical geometry alone. In other words, this asks whether the thickness of the cortical layers can be determined purely from cortical shape, a capability in high demand in both medical image analysis. We expected the *α* obtained through the morphogenic tracks under the EVE assumption to match the measured *α*. As a control, we compared the prediction from the morphogenic tracks against *α* derived from a Laplacian harmonic (LH) potential field, an approach commonly used to establish correspondence between the white matter and pial surfaces (Jones et al., 2000) (**Figure 5D**). We found that the *α* predicted from EVE with morphogenic tracks closely matched the actual *α*, whereas EVE with LH performed worse (**Figure 5E**). These results indicate that the geometry of the cortex alone is sufficient for determining the laminar positions. We further refined the laminar boundary prediction by incorporating the t-constant *α* as prior knowledge with *f*′(*t*) ≈ *t* − 1/2 approximation (**Methods**), showing that such a prior can enable precise laminar segmentation (**Figure 5F**).

### Cell body shape covaries with the physical displacement of cortical folding

We have analyzed cortical anatomy at the laminar scale (mesoscale), but it remains unclear how laminar scale deformation relates to the shape and distribution of cells. Understanding this relationship would allow us to recover the cell density that would exist in the absence of folding-associated deformation. Because this undeformed density reflects the developmental output of the underlying RG cells, it would further enable “retrodictive” inference, such as estimating how many cells were generated per RG cell. This quantity is defined relative to a discrete developmental unit rather than an arbitrarily chosen region, making it straightforward to interpret biologically and providing a common reference for comparing samples. Such framework is particularly valuable for multi-sample pathological comparison of the cerebral cortex using tissue clearing.

We first asked whether cell shape reflects the local tissue deformation caused by cortical folding. Using rRNA staining and ZenCell segmentation (Murakami et al., 2026), we detected the shapes of cell bodies throughout the imaged volume (**Figure 6A**). We confirmed the elongated cell shapes in the deep layers, tangentially longer in sulci and radially stretched in gyri (**Figure 6B**), consistent with Bok’s observation (Bok, 1929). To quantify this anisotropy of cell shape, we calculated the cell lengths along the radial and tangential directions, *ar* and *at* using the directional vector field of the morphogenic tracks (**Figure 6C**). Notably, the ratio *ar*⁄*at*, which is an indicator of anisotropy of cell shape, showed a strong correlation, Pearson’s r being 0.75, 0.68, 0.44, 0.22 for each laminar position, with the stretch rate (**Figure 6D**). This suggests that folding-induced deformation is a physical shape change, with cells deforming in accordance with the laminar scale tissue stretch. This observation, in which laminar scale deformation appears to directly alter cell shape, is consistent with the timing of cortical folding, which occurs after the laminar structure has been largely established (Llinares-Benadero and Borrell, 2019). Because the cells are already in place when folding begins, folding deforms cell shapes accordingly. Building on this understanding, we examined whether we could reconstruct the cell density distribution that would exist in an idealized, fold-free state. Our attempt corresponds to reconstructing the cell distribution as it would appear under unconstrained expansion without folding (**Figure 6E**). This can be achieved through the EVE-based coordinate transformation described above. If the proliferation profile of RG cells is homogeneous within a region, the coordinate transformation yields both the RG cell density distribution *σ*2*D*(*u*, *v*) and the RG cell proliferation profile *ρ*(*t*) (**Figure 6F**). To test this, we first transformed the cell positions, defined as the centroids of the segmented cell bodies, into *uvt* space, and used SVD to assess whether the distribution could be decomposed into *σ*2*D*(*u*, *v*) and *ρ*(*t*). The rank-1 energy fraction was 96.7%, confirming that the distribution is approximately separable into *σ*2*D*(*u*, *v*) and *ρ*(*t*) (**Figure 6G**).

**Figure 6.**
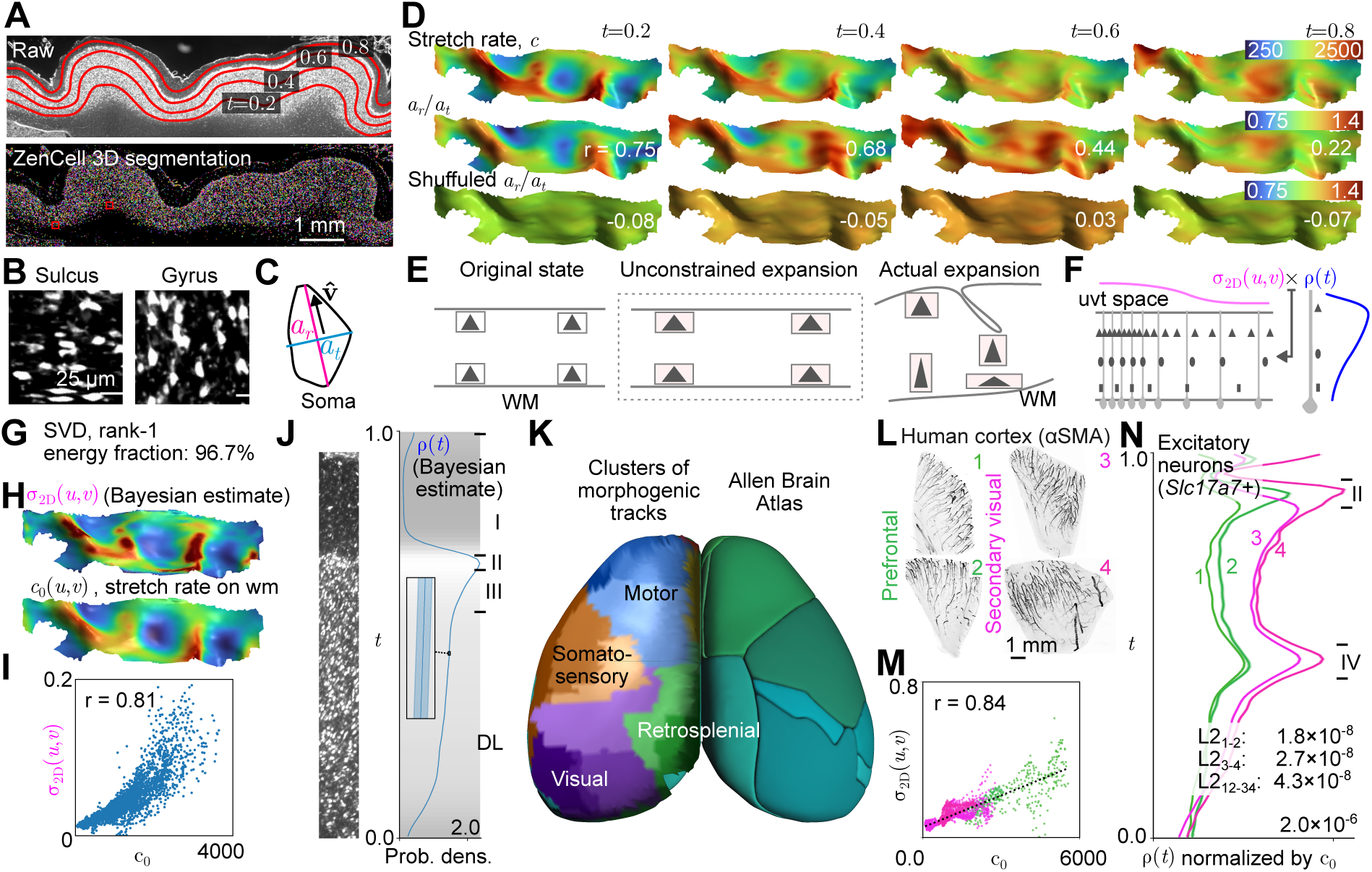
Bayesian retrodictive inference of proliferation rate of RG cells. (**A**) Cell segmentation of the pig cingulate cortex. A representative section of the original image (top) and the segmented cells (bottom) are shown. The boxed areas are shown magnified in **B**. (**B**) Magnified views of rRNA-stained cells in the sulcus and gyrus in a deep layer. (**C**) Definition of *ar* and *at* of a cell. (**D**) Heatmaps of the stretch rate and *ar*⁄*at* and their correlations. Various laminar positions are shown with their Pearson’s r. (**E**) Illustration of the association between cortical folding and cell shape. (**F**) Illustration of the separable cell density functions that collectively describe the cell density distribution. (**G**) SVD rank-1 energy fraction of cell density in *uvt* space. (**H**) Heatmaps of the Bayesian-estimated density function *σ*_2_*D* and *c*_0_. (**I**) Scatter plot of the relationship between *σ*_2_*D* and *c*_0_; Pearson’s r is shown. (**J**) Plot of the Bayesian-estimated density function *ρ*(*t*). The rRNA staining is shown on the left for reference. Error bars show the 95% credible interval. (**K**) Spatial clustering of morphogenic tracks in a mouse brain using cell density around the tracks. The ABA functional boundaries are shown on the right for reference. (**L**) 3D rendering of the four cortical pieces from a human brain with αSMA staining. (**M**) Scatter plot of the relationship between *σ*_2_*D* and *c*_0_; Pearson’s r is shown. (**N**) Plot of the Bayesian-estimated density function *ρ* for each piece.

### Bayesian retrodictive inference of proliferation rate of RG cells

Because the spatial distribution of cells reflects the cumulative outcome of inside-out proliferation, we used the cell position ns to retrodict the proliferation profile of RG cells. Cell proliferation is inherently variable, so the observed density reflects incidental fluctuation as well as genuine developmental change. We therefore adopted a Bayesian approach to quantify the uncertainty in the inferred profile and to distinguish genuine changes in proliferation from incidental fluctuation. Using Bayesian log-Gaussian Cox process (LGCP) inference, we obtained *σ*2*D*(*u*, *v*) and *ρ*(*t*) (**Methods**). *σ*2*D*(*u*, *v*) showed a high correlation with the predicted density of RG fibers, *c*_0_, on white matter surface (**Figure 6H** and **I**). This supports the validity of the *uv* coordinate transformation shown in **Figure 4B**. The *ρ*(*t*) corresponds to the RG cell proliferation profile without the confounding effect of folding (**Figure 6J** and **Methods**), which we expected to be a highly useful metric for comparative analysis. As a demonstration of comparative analysis, we computed *ρ*(*t*) for each morphogenic track in a mouse brain and performed spatial clustering. The cluster assignments changed in accordance with the functional parcellation of the brain (**Figure 6K**), reflecting the fact that radial cell distribution patterns are associated with functional specialization, consistent with Brodmann’s seminal work (Brodmann, 1909).

To test the utility of our workflow for human brain analysis, we performed a regional comparison. We dissected two pieces of prefrontal cortex and two pieces of secondary visual cortex from a single individual (**Figure 6L**). After 3D imaging of excitatory neurons by staining *Slc17a7* mRNA, we quantitatively analyzed how the two areas differ in the absence of folding-induced effects. We confirmed that *σ*2*D*(*u*, *v*) showed a high correlation with the predicted density of RG fibers (r = 0.84, **Figure 6M**), and then calculated the RG cell profiles. The profiles were similar when pieces came from the same region (L2 norm 1.8×10-8 and 2.7×10-8) but distinct when they came from different regions (L2 norm 4.3×10-8, **Figure 6N**), demonstrating the utility of the profile for the comparison purpose.

### A protomap directs cortical folding through the EVE model

So far, we have proposed that RG cell proliferation profiles are uniform within a cortical region (EVE growth model) and that differences between these profiles encode the inter-regional differences of the cortex (Figure 6). Yet one question remains open. If proliferation is uniform, how can folding occur? If all RG cells expanded at the same rate, the naive expectation would be radial growth that preserves the shape of the subventricular zone (protomap), producing a cortex without folds. The obvious explanation would be that RG cells actually grow at different rates in different places, and that this non-uniform growth drives folding. For this to hold, the protomap would have to specify the growth rate at every point precisely enough to reproduce the entire folding pattern. Such fine-grained spatial control seems unlikely, as the protomap is thought to be specified by relatively smooth molecular gradients rather than precise point-by-point instructions (O’Leary et al., 2007). Moreover, if such precise control existed, it would need to be implemented independently in each hemisphere to account for the known left-right differences in folding pattern (Hill et al., 2010), which also seems unlikely. We therefore hypothesized that cortical folding can arise even under the EVE growth model. This view is consistent with the idea that cortical folding arises as a consequence of physical processes. In particular, the differential tangential expansion (DTE) folding theory has elegantly demonstrated, through both simulation and 3D-printed layered gel experiments, that folding can emerge under the assumption of a uniform tangential expansion of the gray matter (Richman et al., 1975; Tallinen et al., 2014, 2016). This assumption of uniform expansion is closely related to equivolume expansion.

In our simulation of DTE, rather than treating the tangential expansion rate as strictly uniform, we allow it to vary in proportion to RG cell density, to test whether folding can arise under the EVE growth model (**Methods**). We first confirmed that our modified DTE produces cortical folding (**Figure 7A**). The simulated stretch rate varied linearly along the radial direction (i.e., t-constant *α*), consistent with our observations (**Figure 7A** and **B**). This supports the interpretation that our observation of t-constant *α* arises as a consequence of folding.

**Figure 7.**
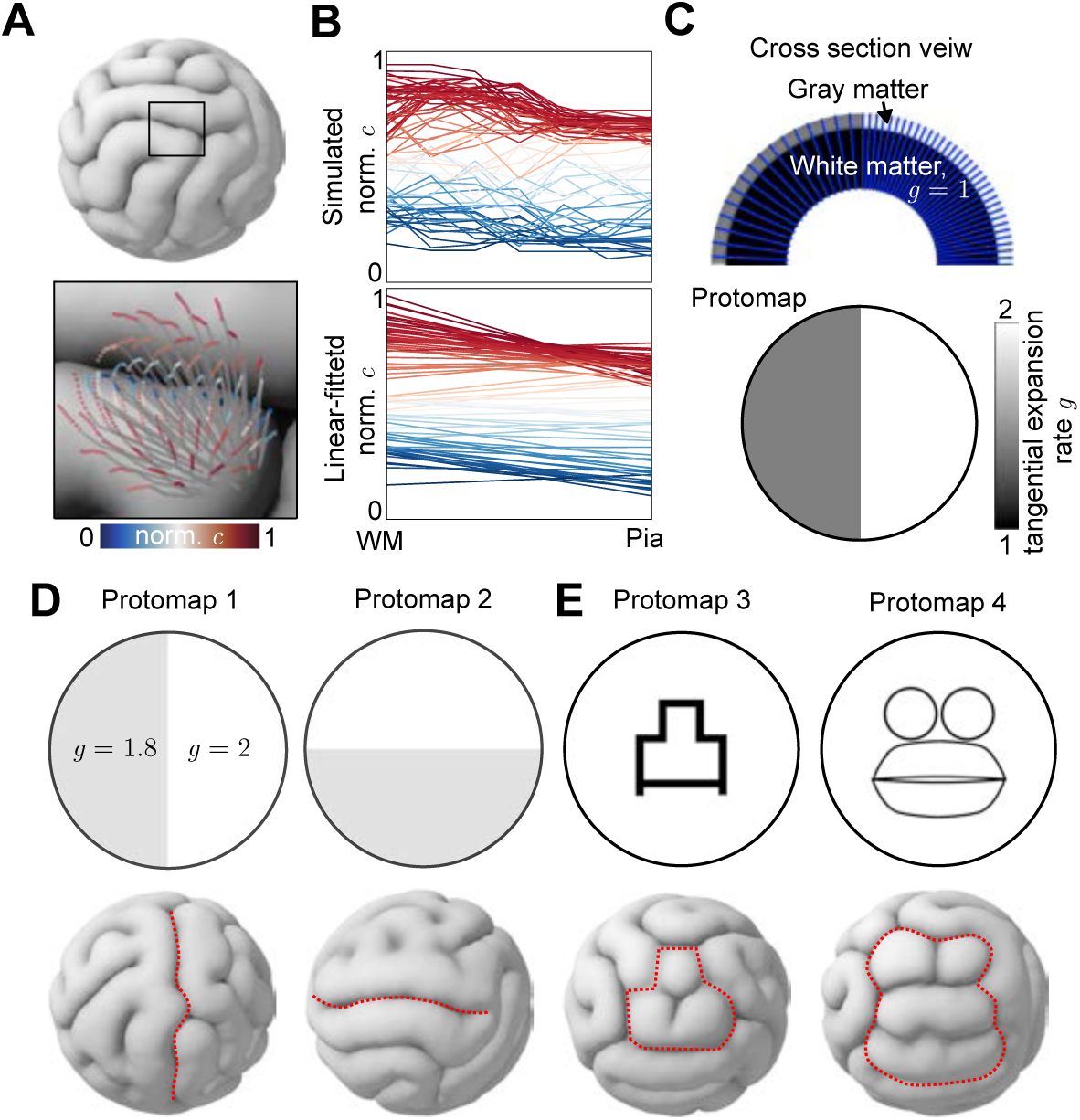
EVE-grounded protomap directs cortical folding within differential tangential expansion theory. (**A**) Simulation result of DTE using a sphere. A magnified view with the simulated morphogenic tracks is shown, with the simulated *c* color-coded at the bottom. (**B**) Plot of simulated *c* from *Swm* to *Spia*. Linearly fitted normalized *c* is shown at the bottom. (**C**) Illustration of protomap-controlled DTE with the EVE assumption. Blue lines indicate RG fibers. The protomap controls the density of RG fibers. (**D**) Protomap-directed cortical folding. The directed sulci are indicated by dotted red lines. (**E**) Protomap-directed cortical folding with a more complex pattern. The directed sulci are indicated by dotted lines.

A limitation of the DTE folding theory is that it cannot easily control the spatial pattern of folding and therefore cannot explain why canonical sulci such as the central sulcus are consistently present despite inter-individual variation in brain shape. We next tested whether the integration of EVE growth model can provide the mechanism that the DTE folding theory lacks for generating canonical sulci. Specifically, we prepared several protomap patterns representing different density distributions of RG cells in the subventricular zone (**Figure 7C**). We then tested whether these protomaps could direct the formation of sulci. We found that a modest reduction in the density of RG cells was sufficient to induce a sulcus at the targeted location (**Figure 7D**). This indicates that even small differences in RG cell densities can give rise to canonical sulci, suggesting that the protomap can specify canonical folding patterns through simple, low-cost density differences rather than precise spatial control of molecular gradient. The manipulation of folding patterns through the protomap was also shown to be relatively flexible, capable of generating a range of patterns (**Figure 7E**). Conventionally, theories governing mesoscale laminar thickness and those governing macroscale folding patterns have existed independently. The EVE growth model offers a framework that can unify these two scales of cortical organization.

## DISCUSSION

In this study, we proposed a new conceptual framework for 3D image analysis based on morphogenic tracks. The central idea is a coordinate transformation grounded in the developmental organization of the cortex. Importantly, this transformation is a tool that facilitates quantification by offering researchers a new perspective without the loss of any information. For example, the original cell density is always exactly recoverable from the transformed space through a simple mathematical operation (multiplying by the Jacobian determinant). The transformation reorganizes spatial information rather than destroying it. The principal advantage of the coordinate transformation is that it makes the tools of mathematical analysis, such as calculus and spatial statistics, readily applicable. It enables analysis at a finer granularity than the conventional binary division into gyri and sulci, allowing deeper investigation across scales, including laminar structure (Figure 5), cell distribution (Figure 6), and the mechanism of folding formation (Figure 7).

We proposed a Bayesian method for inferring the proliferation rate of RG cells, which is particularly valuable for analyzing sparse cell populations. For example, somatostatin-expressing neurons in the human cortex are present in relatively low numbers but are of considerable interest given their implication in neurological disorders such as Alzheimer’s disease (Gabitto et al., 2024). Estimating the density of such sparse populations is subject to large inter-experimenter variability, making pathological comparison difficult. Bayesian inference is particularly powerful in this regime: rather than producing a single unstable estimate from few data points, it yields a full posterior distribution that quantifies the uncertainty of the estimate and naturally regularizes it where data are scarce. This allows genuine biological differences to be distinguished from sampling noise, which is expected to facilitate discovery through comparison.

*limitations of the study*. Because the absolute number of RG fibers is unknown, the inferred proliferation rate per RG cell rests on the assumption that a constant volume is associated with each RG fiber. Although we expect this to hold within the same brain region, we must be aware that “constant volume” may vary depending on the brain regions. The interpretation of the inferred “proliferation rate per RG cell” must be made carefully. Consequently, when comparing quantitative values obtained from different preparations, such as expansion microscopy and conventional cleared samples, normalization with reference to the magnification factor is required. Using the number of penetrating arteries and radial neurites as a proxy could address this scaling challenge, and an image analysis method that accurately counts these structures would open a path to a solution. Another consideration is that our coordinate framework is grounded in RG-derived proliferation, yet some cortical cells, such as inhibitory neurons and oligodendrocyte precursor cells, arrive through tangential migration rather than radial proliferation. The framework still applies to these populations as long as they were uniformly distributed before the onset of folding, since their present distribution would then reflect folding-induced deformation alone. We do not assume this to be the case a priori; rather, our Bayesian point process workflow provides a direct means to test it for each cell type, which we can address through cell-type-specific staining such as mFISH3D (Murakami et al., 2026).

Applying morphogenic tracks to the cerebellum is another intriguing direction. The cerebellum exhibits regular folding at a higher spatial frequency than the cerebral cortex, and whether this can be explained by the same framework remains an open question. Ultimately, morphogenic tracks offer a way to read the history of a tissue’s development from its mature form alone.

## MODEL SYSTEMS AND PERMISSIONS

### Mice Model

C57BL/6J or B6.Cg-Tg(Thy1-YFP)HJrs/J mice from Jackson laboratory were used for the experiments. All procedures involving mice were approved by The Rockefeller University Institutional Animal Care and Use Committee (IACUC) and were in accordance with the National Institutes of Health guidelines.

### Collection of brain

#### Mouse

The 8-week-old mice were sacrificed by an overdose of pentobarbital (> 100 mg/kg), then transcardially perfused with 15 ml of phosphate buffer saline (PBS) and 20 ml of 4% paraformaldehyde (PFA, Electron Microscopy) in PBS. E16.5 embryos were harvested following maternal euthanasia by pentobarbital overdose, followed immediately by brain dissection. The brains were dissected, then post-fixed with 40 mL of 4% PFA in PBS at 4°C overnight before the 3D staining process.

#### Pig

A fresh juvenile pig brain was obtained from a local butcher and dissected. We fixed the tissue with 40 mL of 4% PFA in PBS at 4°C overnight. The fixed tissue was brought to the staining process as in a mouse brain.

#### Human Tissue

Fresh frozen post-mortem human tissue samples were obtained from Science Care. Project approval was sought and granted by The Rockefeller University Institutional Review Board. Tissue donors were de-identified before receipt of tissue. A female donor 65 years of age with no known history of neuropsychiatric or neurological conditions was used in this study. After identification of the prefrontal and secondary visual cortex, the fresh-frozen sample blocks underwent further sub-dissection. After the sub-dissection, we moved the samples to-20°C and incubated for more than 3 hours. We then fixed the tissue with 40 mL of 4% PFA in PBS at 4°C overnight. The fixed tissue was brought to the staining process.

## METHOD DETAILS

### mFISH3D

We performed 3D in situ hybridization and immunostaining according to the published protocol (Murakami et al., 2026). The oligonucleotides and antibodies used in this study are shown in the **Resource Table**. For mouse and pig brains, we performed rRNA staining using mFISH3D. For human brains, we performed Slc17a7 staining. Immunostaining was performed as described in Murakami et al. (2026). When it followed in situ hybridization, the hybridization signals were bleached beforehand (Murakami et al., 2025).

### Microscopy

The Cleared Tissue LightSheet (CTLS) system from Intelligent Imaging Innovations was employed for imaging the cleared tissue samples. The detailed specifications are shown in the previous publication (Murakami et al., 2026). Optical zoom of 10× (human) or 5× (others) was applied, resulting in pixel sizes of 0.65 μm (human) or 1.3 μm (others). A 3 μm (human) or 2 μm (others) z-step was used. All hardware was operated using Slidebook (Intelligent Imaging Innovations), and data was stored on a Dell PowerEdge R740XD with twelve 16 TB hard drives with RAID configuration via a 10-GB Ethernet fiber network.

### Image Analysis

#### Stitching, correcting chromatic aberration, and image registration

Stitching and correction of chromatic aberrations were performed as described previously using BigStitcher (Murakami et al., 2026). For the mouse brain, we registered the autofluorescence images to the Allen Brain Atlas. When necessary, we also registered images across multiple rounds, as previously reported (Murakami et al., 2026).

#### Cell segmentation

We used ZenCell for cell segmentation (Murakami et al., 2026). Otherwise, we used a published dataset (Figure 6K and 6L–N) (Murakami et al., 2018; Murakami and Heintz, 2022).

### Morphogenic track analysis

#### Surface mesh construction

The pial and WM surfaces were reconstructed from manually drawn contours using a custom Napari plugin (napari-spline) followed by Poisson surface reconstruction. On evenly spaced *z*-planes spanning the structure of interest, contours were drawn interactively along the target boundary using the napari-spline plugin. Each contour is represented as a piecewise cubic Bézier path. Each Bézier path was resampled to 20 points spaced evenly along its arc length. The resampled contours were concatenated into a single 3D point cloud. A triangular mesh was reconstructed from the aggregated point cloud using screened Poisson surface reconstruction. If necessary, reconstructed surfaces were manually corrected using Blender. Each mesh was then uniformly resampled to a prescribed triangle density using the approximate centroidal Voronoi diagram algorithm (Valette and Chassery, 2004). Both the pial and white-matter surfaces were resampled independently with this pipeline and saved in PLY format for downstream analysis. The napari-spline is available at GitHub (https://github.com/tatz-murakami/napari-spline).

#### Extraction of ground-truth directional vectors

The targeted structures were manually traced using the BigTrace Fiji plugin (Katrukha et al., 2025). For penetrating arteries, structures with a clear major branch were selected and traced. For the radial glial fibers, Bergmann glial fibers, and radial neurites in Figure 2, thick fibers were selected and traced. Unit directional vectors were extracted at the starting point, anchor points, and end point of each track and used for downstream analysis.

### Prediction of the unit directional vector field

#### Regression model and data

The unit directional vector field is parameterized by a multi-layer perceptron (MLP) **v***_θ_*(**x**) with parameters *θ*, and the unit field used downstream is obtained by l_2_ - normalisation, v̂*_θ_*(**x**) = **v***_θ_*(**x**)/∥ **v***_θ_*(**x**) ∥_2_. Inputs are normalised to the unit cube x̃ = (**x** − **x**_min_)/(**x**_max_ − **x**_min_), where **x**_min_, **x**_max_are the joint axis-aligned bounding-box extrema of the pial and white-matter meshes. Supervision was provided by a set of *N* orientation targets {(**x**, **u**)}*^N^*^obs^, where 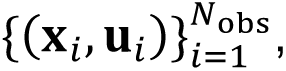 is an location and 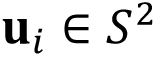 is the local unit orientation extracted at that location. For the interior regularization terms, collocation points {**x**^int^} were sampled uniformly at random inside the joint bounding box at every optimization step.

#### Loss functions

The training objective combines three terms,

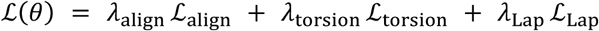

with weights (*λ*_align_, *λ*_torsion_, *λ*_Lap_) = (1, 10^−6^, 10^−2^) chosen empirically so that the three terms operate on comparable scales during training.

*(i) Orientation alignment.* We adopted a cosine-similarity that penalizes the deviation of the predicted unit field from the observed local orientation at each point:

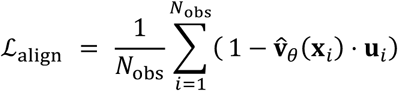

*(ii) Curvature-weighted torsion regularization.* We found that jointly regularizing curvature and torsion effectively prevents overfitting to noise in the orientation targets. The joint penalty is implemented via the triple product **T** ⋅ (**T**′ × **T**″), which by the Frenet–Serret relations equals *κ*^2^*τ* for arclength-parameterised curves, where *κ* is the curvature and *τ* the torsion. Its square therefore penalizes both quantities simultaneously. For each collocation point **x**_0_ = **x**^int^ we perform two short explicit-Euler steps of length *ε* along the predicted unit field, **x***_k_*_+1_ = **x***_k_* + *ε* **T***_k_* with **T***_k_* = v̂*_θ_*(**x***_k_*) for *k* = 0,1, yielding the three tangents **T**_0_, **T**_1_, **T**_2_. Central finite differences give the discrete arclength derivatives **T**′ ≈ (**T**_2_ − **T**_0_)/(2*ε*) and **T**″ ≈ (**T**_2_ − 2**T**_1_ + **T**_0_)/*ε*^2^. The loss is

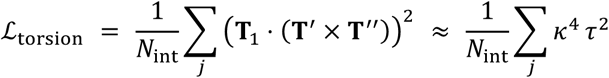

This combined penalty was empirically more robust against overfitting than the curvature-only or torsion-only form. We used *ε* = 10^−3^ in normalised coordinates.

*(iii) Dirichlet (harmonic) regularize.* To discourage sharp spatial variations in the field, which can lead to self-intersecting trajectories, we penalized the Dirichlet energy over the interior collocation points.

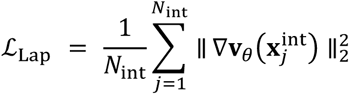

Since minimizers of the Dirichlet energy under fixed boundary data are harmonic, this term drives **v***_θ_* toward a smooth, near-harmonic field. We penalized the Dirichlet energy rather than the Laplacian residual itself, as the latter requires second-order network derivatives that are considerably more expensive to compute. The training was implemented in PyTorch and is available at GitHub.

#### Quantification of directional co-alignment

Co-alignment of penetrating arteries with other radial structures was quantified as the mean angular distance between the model prediction from arteries and the unit directional vectors of the targeted radial structure (Figure 2). For a track containing *N* sample points with reference unit vectors 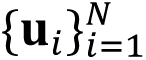and model predictions 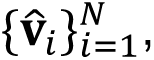 we define *d*_ang_ as the mean angle between prediction and reference, normalised by *π*:

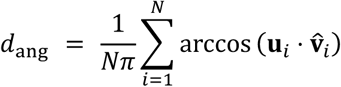

#### Morphogenic track simulation

The simulation of the morphogenic track was done for the visualization purpose. Morphogenic tracks were computed by numerically integrating the ordinary differential equation

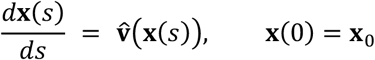

using a fixed-step fourth-order Runge–Kutta scheme. The arclength parameter *s* is expressed in micrometers. Each streamline was then truncated at its first intersection with the pial surface mesh.

### EVE-based coordinate transformation

#### t mapping

To assign an equivolume time coordinate *t* to an arbitrary query point **x***_p_* in the cortical volume, we first determined the entry point **x**_0_ ∈ *S*_wm_ on the white-matter surface and the total trajectory length (cortical thickness) *L*(**x**_0_) along the trained unit field v̂ that passes through **x***_p_*. The condition ∇ ⋅ **v** = 0 is the standard volume-preservation (incompressibility) condition, which is the equivolume principle. Using the stretch rate *c*, the equivolume growth model can be written as

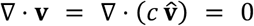

Expanding the divergence with the product rule gives

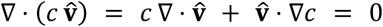

Along the trajectory, *c* becomes a function of *s* via *c*(*s*): = *c*(**x**(*s*)). By the chain rule,

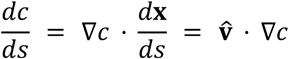

where the second equality uses *d***x**/*ds* = v̂. This yields the ordinary differential equation

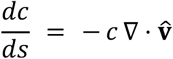

Solving the differential equation by separation of variables with the initial condition

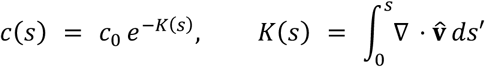

was fixed by imposing unit transit time, ∫*L*(**x**0) *d s*/*c*(*s*) = 1. This gives

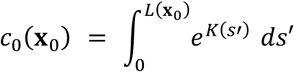

Combining these yields the target equivolume time coordinate

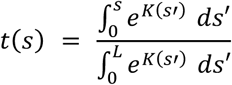

#### Correction of t

A limitation of the divergence-based construction above is its numerical stability. The divergence ∇ ⋅ v̂ can change sharply at sulcal walls, where neighboring trajectories bend in opposite directions and v̂ effectively switches sign across a thin transition. Because the computation of *t* involves repeated integration and exponentiation of the divergence, such local sharp changes can propagate into and distort the reconstructed time coordinate. To stabilize *t*, we imposed a smooth linear-in-*t* prior on the stretch rate *c* and recomputed the time coordinate under this prior. The prior takes the form

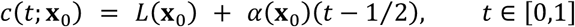

where *α* is the stretch acceleration. This form was justified empirically (Figure 5). A direct linear fit to *c*(*t*; **x**_0_) performed poorly in practice: the fitted profiles were sensitive to noise in the raw *c*(*t*) when *c* was strongly skewed toward one boundary. Fitting in log space avoided the issue and is better justified theoretically: the closed-form *c*(*t*) is exponential in the accumulated divergence *K*(*s*), so log*c* is intrinsically a more natural target for a linear model than *c* itself. Rewriting the formula as

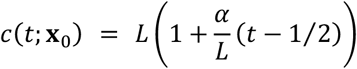

and taking the logarithm gives

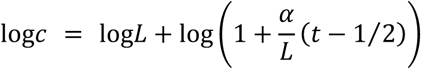

Applying the first-order expansion log(1 + *x*) ≈ *x* for |*x*| < 1 with *x* = (*α*/*L*)(*t* − 1⁄2) yields

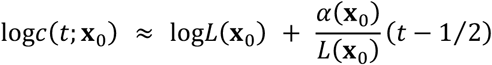

The slope *α*(**x**_0_) was estimated by ordinary least-squares regression of log*c*(*t*; **x**_0_) against (*t* − 1⁄2), and rescaled by *L*(**x**_0_). Prior to fitting, *c*(*t*; **x**_0_) was smoothed across the white-matter surface with umbrella-Laplacian smoothing using the mesh vertex adjacency, to suppress vertex-scale noise before the log transform. The time coordinate *t* was then recomputed using this linearized *c*(*t*; **x**_0_).

#### uv mapping

Points **x**_0_ ∈ *S*_wm_ were mapped into planar (*u*, *v*) coordinates by metric multidimensional scaling (MDS) applied to pairwise shortest-path distances on the equivolumeweighted mesh graph. Each mesh edge (*i*, *j*) was assigned the equivolume-weighted length

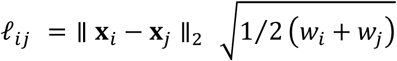

where *w_i_* = *c*_0_ is the per-vertex equivolume weight, so that regions where trajectories emerge more rapidly (larger *c*_0_) receive proportionally larger area in the (*u*, *v*) plane. This yields a bidirectional map **x**_0_ ↔ (*u*, *v*).

### Cell density matrix in a normalized arclength coordinate

The cell density matrix was prepared for the cortical flattening. For each query point

**x***_p_* (segmented cell centroid), we defined a normalized arclength coordinate

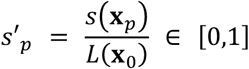

measuring the fractional distance from the white-matter surface to the pial surface along the v̂-trajectory through **x***_p_*. Combined with the UV coordinate (*u_p_*, *v_p_*) of **x***_p_*, the samples {(*u_p_*, *v_p_*, *s*′*_p_*)} were aggregated into a two-dimensional intensity matrix **M** ∈ ℝ*^T^*^×*N*^, whose rows index *T* uniform arclength bins on [0,1] and whose columns index the *N* vertices of the flattened WM mesh. Each entry is the smoothed cell density at that bin and vertex.

### Cortical flattening assuming linearity of c in t

Using the cell density matrix **M**, we fitted the stretch acceleration *α*(*u*, *v*) per vertex by requiring the density profiles at all vertices to become consistent under *c*(*t*) = *L*(*u*, *v*) + *α*(*u*, *v*) (*t* − 1⁄2).

#### Regression model

*α*(*u*, *v*) was parameterized by a neural network consisting of a positional encoding followed by an MLP, with a final bounded activation enforcing |*α*| < 2*L*.

#### Loss

Each vertex column of **M** was first normalized to unit sum. At every optimization step, columns were resampled by linear interpolation at the warped positions obtained by *α*, yielding a warped intensity matrix M̃. This was Gaussian-smoothed along the *t*-axis to suppress high-frequency noise, after which the training loss was

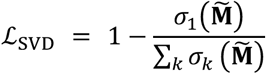

where *σ_k_* are the singular values of M̃. This loss vanishes when M̃ is rank-one — i.e. when every vertex column is a scalar multiple of a common profile — so minimizing it drives all vertices to share a single density profile after reparameterization. The training was implemented in PyTorch and is available at GitHub.

### Cortical flattening without assuming linearity of c in t

To understand the spatial pattern of the stretch rate, we also implemented cortical flattening without assuming linearity of *c*(*t*). Instead of fitting *α*, we fitted *t̃*(*t*; *u*, *v*) (the warped time coordinate) per vertex by requiring the density profiles at all vertices to become consistent under the warp.

#### Regression model

The warp was parameterized by its derivative τ̇(*t*; *u*, *v*) > 0, guaranteeing monotonicity of *t̃* by construction. A neural network output the positive derivative curve, and *t̃*(*t*; *u*, *v*) was recovered by cumulative integration of τ̇ and normalization so that *t̃*(0) = 0 and *t̃*(1) = 1.

#### Loss

The SVD loss ℒ_SVD_ was augmented with a Jacobian regularization on a logarithmic scale:

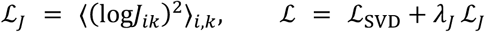

where *J_ik_* is the local Jacobian at vertex *i* and time index *k*. This prevents degenerate warps that collapse or over-stretch *t*. The training was implemented in PyTorch and is available at GitHub.

### Rank-1 energy fraction

To assess whether the spatial patterns of the stretch rate (Figure 5) and the cell density (Figure 6) can be factored as a product of a (*u*, *v*)-only function and a *t*-only function, we computed the rank-1 energy fraction 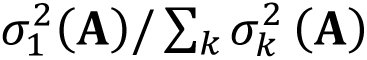 — the fraction of spectral energy captured by the leading singular component — of the corresponding data matrix **A**. Values are reported as percentages.

### Cell shape analysis

Because the cell shapes obtained from segmentation can be affected by the anisotropic point spread function of light-sheet microscopy, we performed the analysis excluding the axial (*z*) dimension, reducing this artifact. To quantify the elongation of each cortical cell relative to the local morphogenic direction, we computed the cell’s extent along a radial direction (aligned with the local flow) and an orthogonal tangential direction in the imaging (*x*, *y*) plane. The projection onto the (*x*, *y*) plane avoided distortion from the non-isotropic resolution along the *z*-axis. At each cell centroid **x***_p_*, the trained unit vector field v̂(**x***_p_*) was projected onto the (*x*, *y*) plane, giving the local radial direction. The local tangential direction was defined as its 90^∘^in-plane rotation. For each cell, the 3-D segmentation mask was max-projected along the *z*-axis to yield a 2-D silhouette. The radial extent *a_r_* and tangential extent *a_t_* were the peak-to-peak ranges of the silhouette pixels projected onto the radial and tangential directions, respectively, and the ratio was computed as *a_r_*/*a_t_*.

### Spatial clustering

For spatial clustering shown in Figure 6K, we used agglomerative hierarchical clustering with Ward linkage, subject to a spatial-connectivity constraint that restricts merges to adjacent units.

### Bayesian point process

We modeled the spatial and temporal density of cortical cells with log-Cox Gaussian process (LCGP) point-process models: the log-rate of a Poisson observation model is decomposed into slowly-varying Gaussian process (GP) components over the flattened UV plane and along the time coordinate *t*. Cells were binned by their (*u*, *v*, *t*) coordinates into a count matrix **Y**, where *Y_ik_* is the number of cells at face *i* within the

*k*-th time bin on [0,1].

#### Marginal temporal density ρ(t)

Summing counts over faces gives the temporal count vector *y_k_^t^* = ∑*_i_ Y_ik_*, modeled as

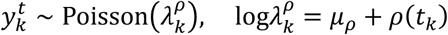

with a Gaussian process prior on *ρ*, approximated via a Hilbert-space Gaussian process (HSGP). The posterior temporal intensity *λ_k_^ρ^* was normalized by the equivolume-weighted area summed over the analyzed faces, so that the reported *ρ*(*t*) reflects a per-unit-volume cell density rather than a probability.

#### Marginal spatial density σ_2D_(u, v)

Summing counts over time gives the face-marginal count vector 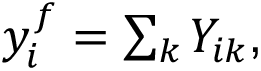 modeled as

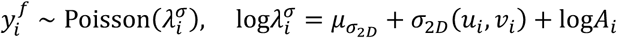

where *A_i_* is the physical 3D surface area of face *i* — included as an offset so that *σ*_2*D*_ is the log-density per unit surface area on the original cortex rather than on the flattened UV plane — and *σ*_2*D*_ carries a 2-D Gaussian process prior, again approximated via HSGP.

#### Simulation of cortical folding

The mechanical model follows Tallinen et al.: a bilayer Neo-Hookean solid comprising a thin cortex bonded to a core, grown under a multiplicatively decomposed de-formation gradient **F** = **F***_e_***F***_g_*, with equal shear modulus in both layers and bulk modulus *K* = 5*μ*. Growth is purely tangential,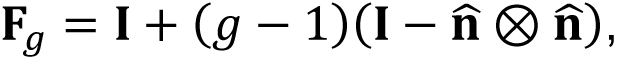 where n̂ is the outward pial normal. The domain is a thick spherical shell, *R*/2 ≤ *r* ≤ *R*, with the inner surface clamped; quasistatic equilibrium is reached by explicit dynamic relaxation with a self-contact penalty, exactly as in the reference. We reimplemented the finite-element solver from scratch in Python (NumPy/Numba, with a CuPy GPU backend) on an irregular tetrahedral mesh resolving the cortex with at least eight element layers. We depart from the reference in the specification of the growth factor *g*. We represent each radial fiber as a radial column of elements and each element as one *segment* of that fiber. Under the equivolume postulate, every segment adds the same absolute tissue volume *Δa* per growth step, giving the per-element increment

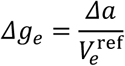

where *V*^ref^ is the reference (undeformed) element volume. The protomap is thereby encoded geometrically: denser fibers pack into narrower columns of smaller reference volume, which by the equation grow fastest and bias the location of emerging sulci and gyri. The script is available at GitHub (https://github.com/tatz-murakami/folding-simulation).

## QUANTIFICATION AND STATISTICAL ANALYSIS

### Statistical Analysis

Statistical analyses were performed by SciPy. The Mann–Whitney U test was used with Bonferroni correction for multiple comparisons. In this study, p < 0.05 was considered as significant.

### Correlation analysis

The linear association between two continuous variables *X* and *Y* was quantified by the Pearson correlation coefficient

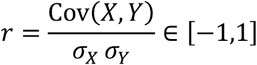

Correlations were computed using SciPy.

#### *L*^2^ distance

The *L*^2^ distance between two functions *f* and *g* was computed as

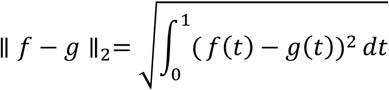

#### Mean squared error

The mean squared error between two vectors **x**, **y** was computed as

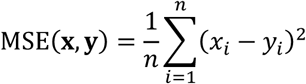

## DATA AND SOFTWARE AVAILABILITY

Our Python codes for the analysis is available on GitHub (https://github.com/tatz-murakami/morphotrack, https://github.com/tatz-murakami/napari-spline, and https://github.com/tatz-murakami/folding-simulation).

## RESOURCE TABLE

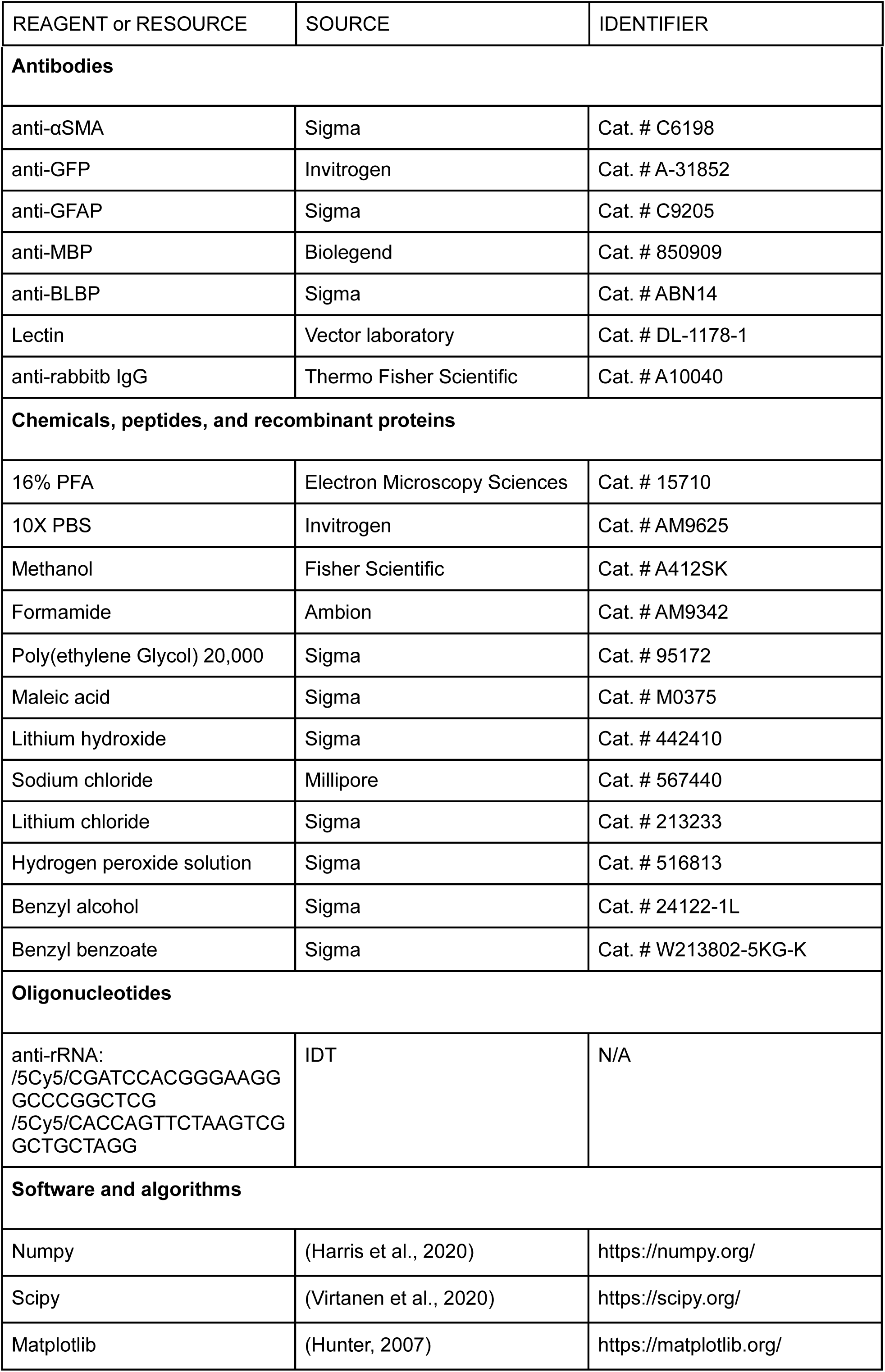
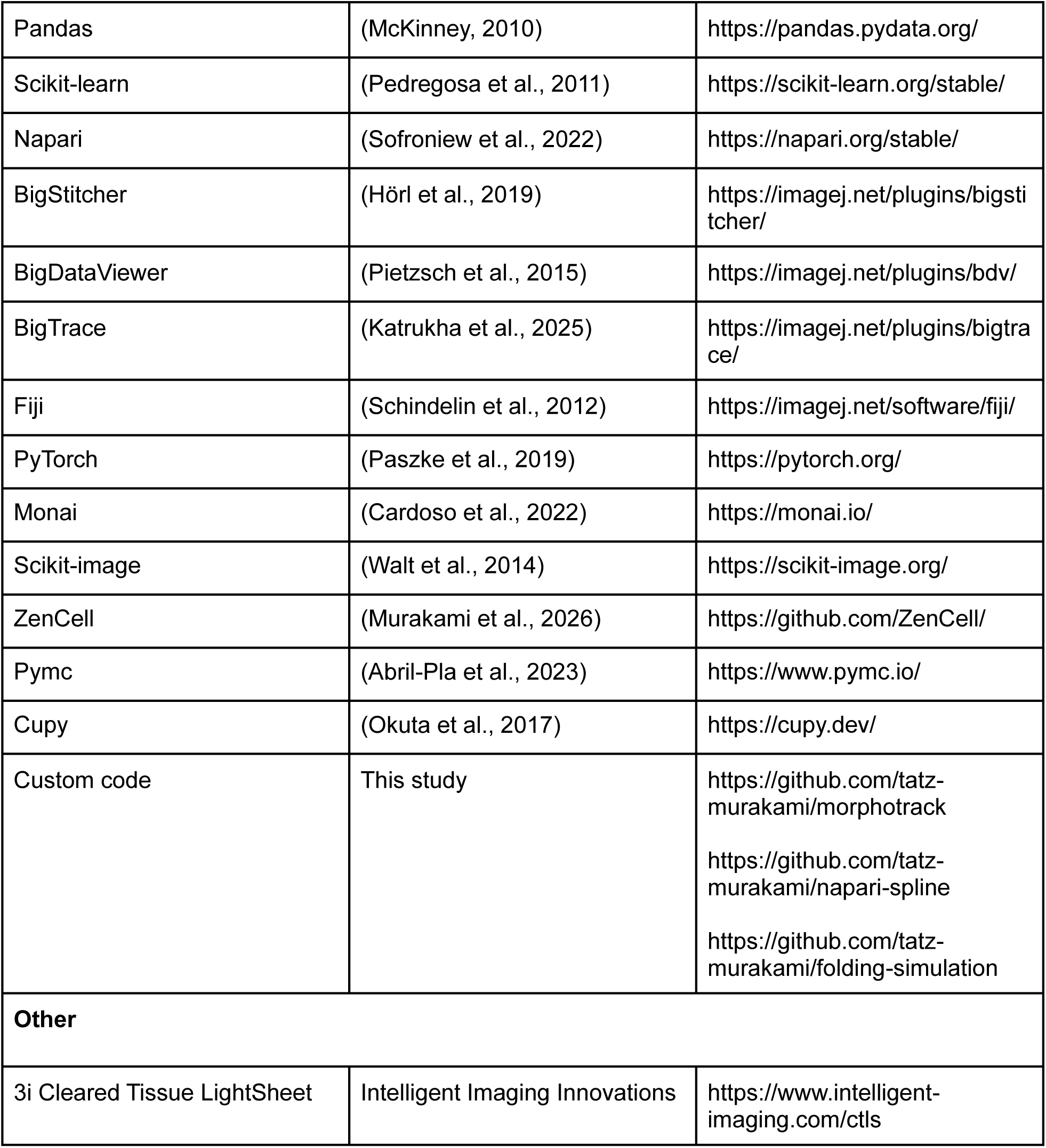

## ACKNOWLEDGEMENTS

This work was supported by CHDI Foundation (to N.H.), Fisher Center for Alzheimer’s Research Foundation (to N.H.), and Leon Levy Scholarships in Neuroscience (to T.C.M.). We thank Yurie Maeda and Jie Xing for their technical assistance in the experiment. The human tissues presented in this study were provided by Miami’s Endowment Brain Bank. The mice were bred in Hatten lab at Rockefeller University and provided by Yin Fang and Zack Schapire. Yuchen Zhang helped harvesting a pig brain. We thank Meng Xia, Ziyi Lin, Yuejia Yin, and Anthony Carvalloza for their support in cell segmentation.

## AUTHOR CONTRIBUTIONS

T.C.M. designed the study. T.C.M. performed the experiments. T.C.M and N.H. supervised the research. T.C.M drafted the manuscript. T.C.M and N.H. reviewed and edited the manuscript. N.H. acquired research funding.

## DECLARATION OF GENERATIVE AI AND AI-ASSISTED TECHNOLOGIES IN THE WRITING PROCESS

During the preparation of this work, the authors used Claude Code Opus 4.7 in order to write Python code and generate associated Materials and Methods. After using this tool/service, the authors reviewed and edited the content as needed and takes full responsibility for the content of the published article.]

## LEAD CONTACT

Further information and requests for resources and reagents should be directed to and will be fulfilled by the Lead Contact.

## SUPPLEMENTARY FIGURE TITLES AND LEGENDS

**Figure S1.**
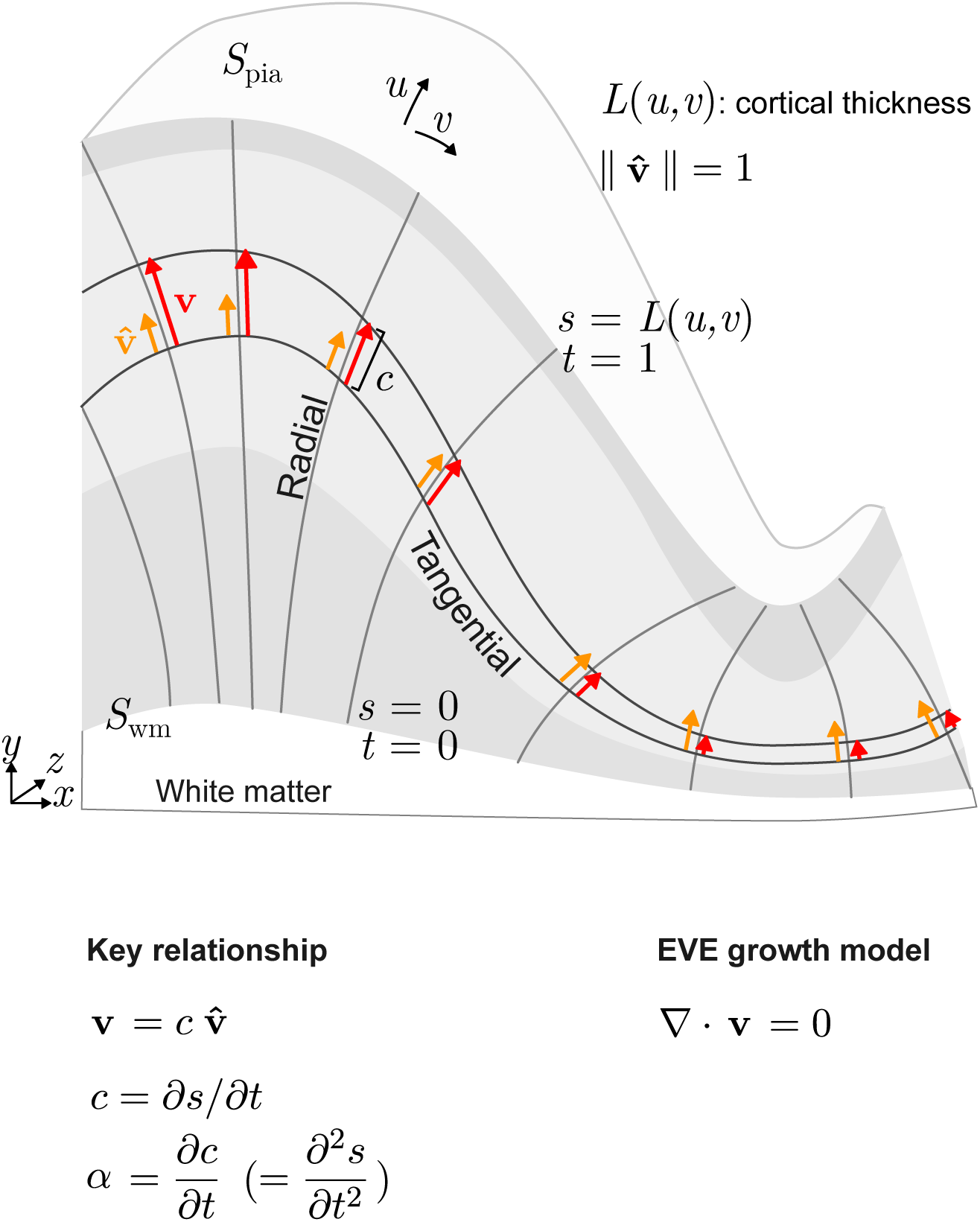
**Geometry and notation of the morphogenic coordinate system.**

## SUPPLEMENTARY VIDEO INFORMATION

**Video S1.** Morphogenic tracks of a mouse brain overlaid with the Allen Brain Atlas. The αSMA signal is also shown on the right.

**Video S2.** Morphogenic tracks of a pig cingulate cortex overlaid with αSMA. The pial vasculature was removed for visibility.

